# Reactive oxygen species generated by infrared laser light in optical tweezers inhibits the germination of bacterial spores

**DOI:** 10.1101/2022.03.29.486222

**Authors:** Dmitry Malyshev, Nicholas Finlay Robinson, Rasmus Öberg, Tobias Dahlberg, Magnus Andersson

## Abstract

Bacterial spores are highly resistant to heat, radiation, and various disinfection chemicals. These impacts on the biophysical and physicochemical properties of spores can be studied on the single-cell level using optical tweezers. However, the effect of the trapping laser on spores’ germination rate is not fully understood. In this work, we assess the impact of 1064 nm laser light on the germination of *Bacillus thuringiensis* spores. The results show that the germination rate of spores after laser exposure follows a sigmoid dose-response relationship, with only 15 % of spores germinating after 20 J of laser light. Under anaerobic growth conditions, the percentage of germinating spores at 20 J increased to 65 %. The results thereby indicate that molecular oxygen is a major contributor to the germination-inhibiting effect observed. Thus, our study highlights the risk for optical trapping of spores and ways to mitigate it.

## 1 INTRODUCTION

Bacterial spores are a dormant form of bacteria exhibiting no cellular activity. They are highly resilient, able to survive for years in natural conditions, as well as capable of surviving a number of decontamination methods ^1,2^. Due to their resilience, pathogenic spores are a hazard in healthcare alongside food production and storage, with some bacterial spores like the anthrax-causing *B. anthracis* being classed as biological warfare agents ^3,4^. It is therefore important to better understand their resilience to develop robust decontamination methods to be able to diagnose and detect spores. However, spores of *Bacillus* and *Clostridium* species exhibit significant heterogeneity, in which individual spores show great variation in germination rate, metabolic activity, and conditions for heat activation before germination ^5,6,7^. Therefore, tools that can study these mechanisms on an individual spore level are needed since the heterogeneity is masked in bulk studies.

Optical tweezers are versatile tools that can measure the biophysical and physicochemical properties of individual spores. For example, optical tweezers can be used to measure adhesion forces, hydrodynamic coefficients, and Raman scattering from individual bacteria/spores ^8,9,10^. An optical tweezers system focuses a laser beam down to a sub-micrometer spot, generating an attractive force sufficient to trap and hold micro-sized objects. Laser trapping of biological objects has been considered largely non-invasive using near-infrared lasers (NIR) lasers at low laser powers, order of mW ^11^. However, it has been shown that even laser traps with powers as low as 3 mW ^12,13^, and doses as low as 0.54 J ^14^ can affect cell viability. Previous studies suggest that intense laser irradiation may inflict DNA damage in cells, as well as produce reactive oxygen species (ROS). Particularly interesting is the generation of singlet oxygen ^15,16,17^, which in turn affects the function and structural integrity of the cells ^12,13,18^. However, compared to cells, spores are significantly more resilient to thermal and radiation damage from optical tweezers since they have several mechanisms to protect their DNA.

Optical tweezers have been used extensively to characterize and assess spore mechanisms on an individual level ^19,20,21,22,9,23,24,25^, but the number of studies detailing the impact laser trapping has on the spore is limited. In a recent work it was shown that optical tweezers could cause structural changes at high dose levels. It was also shown that laser light can significantly enhance chemical reactions in spores, such as spore degradation by sodium hypochlorite ^10^. However, the mechanism responsible for this degradation was not clearly identified.

In this work we explore how the germination rate is affected by irradiating bacterial spores with a 1064 nm laser beam, a common laser wavelength used in optical tweezers. Based on previous literature, a 1064 nm wavelength should generate a significant amount of ROS from dissolved oxygen ^26^. Therefore, we exposed spores in a liquid growth media to various radiation doses and observed the subsequent germination process, seen as the increase in physical size. A decrease in the number of spores germinated in comparison to a control would indicate chemical changes in the spores from laser exposure. To distinguish whether damage from light exposure is due to the generation of ROS, we further expose anaerobic spore samples to the same dose and observe whether the number of damaged spores is similar to that of the aerobic spores.

## 2 EXPERIMENTAL METHODS

### 2.1 Laser system and microscope

The laser system used is part of the optical tweezers (laser tweezers) Raman Spectroscopy system previously described in ^27,28,29^. Briefly, a Gaussian laser beam operating at 1064 nm is coupled into the microscope using a dichroic shortpass mirror with a cut off wavelength of 650 nm. Imaging and focusing of the beam is done by a 60× water immersion objective (UPlanSApo60, Olympus) with a numerical aperture of 1.2 and a working distance of 0.28 mm. This provides a diffraction-limited spot diameter in the focal plane of ∼ 1 *µ*m. The setup has been build to have low drifts keeping both sample temperature, focal position, and imaging conditions stable for long periods of time, ∼ 1 several hours.

### 2.2 Sample preparation

*Bacillus thuringiensis* ATCC 12435 cells were grown on BBLK agar (BD) plates, incubated at 30 °C overnight. Cells were collected by scraping them off the agar and transferred to a 1.5 ml Eppendorf tube and centrifuged once to remove leftover growth media. To allow sporulation, the cells were then stored at 4 °C overnight.

The resulting spore suspension was then purified by centrifuging in deionised water five times, discarding the supernatant and resuspending the pellet each time. After being purified, the culture was resuspended in deionised water and stored at 4 °C.

### 2.3 Sample preparation and exposing spores to laser light

A sample was made by adding a 1 cm diameter ring of 1 mm thick vacuum grease on a 24 × 60 mm glass coverslip (no 1, Paul Marienfeld GmbH & Co). 2 *µ*L of purified spore suspension was placed inside the ring and left to dry. Then, 10 *µ*L of TSB broth was added on top of the dried sample and sealed with a 23 mm × 23 mm glass coverslip for observation under the microscope. For anaerobic experiments, the TSB broth was degassed overnight in a 2.5 l anaerobic jar (Oxoid), using anaerobic sachets (AnaeroGen 2.5 l, Oxoid). The TSB and dried sample were then placed inside a nitrogen flushed glovebox for 15 min to ensure oxygen displacement from the box. The broth was then added and the sample sealed. The rest of the experiment was carried out in the same way as the aerobic experiments.

To irradiate spores, we first determined the focal position of the laser beam by trapping a non-settled spore. The position of the trapped spore clearly indicates the focal volume. This position was registered in a xy-coordinate system using a in-house LabView program. Spores settled on the cover slide were then illuminated using different dose levels (power × time). We used 1 J (1.0 W, 1 sec), 10 J (1.0 W, 10 sec), 20 J (1.0 W, 20 sec) and 50 J (1.0 W, 50 sec).

After irradiating the spores, we heated the objective nosecone to 30 °C to start spore germination. We observed the spores over a period of 120 minutes, taking an image of the field of view every 10 minutes. For each power setting, we imaged at least 80 individual spores (technical replicates), over at least 3 separate experiments (biological replicates). We used non-exposed spores as controls.

### 2.4 SEM microscopy

To perform SEM imaging, the spore incubation was carried out as detailed above, with two modifications. We replaced the grease ring with a PDMS ring to more easily open the seal. Furthermore, we stopped the spore incubation after 60 min instead of 120 min, as after 2 hours many germinated spores were lost from the slide during the washing process.

After 1 hour of incubation, the sample was gently rinsed with 70 % ethanol, followed by 100 % ethanol. We then coated the sample with a ∼5 nm layer of platinum using a Quorum Q150T-ES sputter coater. We imaged samples using a Carl Zeiss Merlin FESEM electron microscope using the InLens imaging mode at a magnification of 15,000×.

### 2.5 Data Analysis

To determine whether a spore was in the process of germinating or not, we observed the morphological growth of the spore. The physical size of the spore increases as the spore germinates and grows into a vegetative cell. We measured the spore size as the area occupied by the spores in the previously mentioned field of view images. To analyze the images with a degree of automation, we used the area selection tools of ImageJ Fiji 1.53e ^30^. The procedure used to obtain automated measurements is listed in the supporting information. We consider the spores germinating if they during incubation grow more than 50 % from their initial size.

Statistical analysis was done using Graphpad Prism 9. We used Kruskal-Wallis test with Dunn’s multiple comparisons to determine the statistical significance of the changes in spore outgrowth. To compare binary outcomes (germination or lack of germination), we used Fisher’s exact tests. We used the Wilson/Brown method to compute confidence intervals for the percentage of germinating spores, and all curves were fitted in Origin 2018.

## 3 RESULTS AND DISCUSSION

Optical tweezers allow us to study trapped single spores’ biophysical and physicochemical properties over longer periods of time. NIR-lasers are often used for trapping since measurements performed in water or buffer solutions have low absorption in this wavelength region. Despite low absorption, the intensity of the laser beam is very high since a high-numerical objective is used to focus the beam into a diffraction-limited spot ^31^. The total irradiation dose can therefore affect spores through phototoxic effects, which are well summarised for bacterial cells in ^26^. Compared to bacterial cells, spores are extremely resilient to environmental stresses, but they are also metabolically inactive and therefore limited in their ability to respond to ROS. To investigate the influence of ROS on spores’ germination rate, we first looked at how the germination rate of laser-irradiated spores exposed to various laser doses compares to that of non-irradiated spores.

It should be noted that lack of germination after laser exposure does not necessarily mean the spores are dead. Spores exposed to laser light may grow slower or have delayed germination. Therefore, longer exposure time and removing any dividing cells would be needed to investigate this further. However, since spores were given 2 h to germinate in favorable conditions (high nutrition levels and 30 °C,), most likely non-germinating spores were significantly damaged.

At low doses (1 J), spores in TSB germinated into vegetative cells within 2 hours, with no difference seen between the exposed and non-exposed parts of the field of view, see Figure 1 A. By contrast, when spores are exposed to 50 J, most spores fail to germinate, while the non-exposed spores in the same field of view germinate normally, see Figure 1 B. Some spores have even lost their stored CaDPA, as seen from the change in refractive index, indicating that the high irradiation dose might have damaged the spore body. The inhibited germination is in line with the prediction that generated ROS will react with and damage DNA as well important cellular machinery ^32^. ROS are involved in base excision and moderately involved in single-strand DNA breaks and thymine dimerization ^33,34,35^. It has been speculated that the lower hydration level of the spore core might offer some protection from ROS ^36^. However, since the core is partially hydrated ^37^, ROS can likely be generated directly inside the core, bypassing the protective outer layers of the spore, such as the coat, cortex, and membrane.

To further decipher ROS impact on germination rate we quantified the percentage of spores growing into vegetative cells (germinating) after exposure to different doses of irradiation, with examples of the changes in the field of view shown in Figures S1 - S3. The percentage of germinated cells decreased from 87 % (n=112) in the unexposed control, to 83 % (n=94) with 1 J exposure, 76 % (n=83) with 10 J, 15 % (n=94) with 20 J, and 7% (n=91) with 50 J, see Figure 1 C. We conclude that 20 J and 50 J laser irradiation had a statistically significant effect on spore germination compared to the control (p *<* 0.0001 for both). For spores that were exposed to 10 J, no statistically significant effect was seen (p = 0.09), and no difference was seen between control and 1 J (p = 0.64).

**FIGURE 1.**
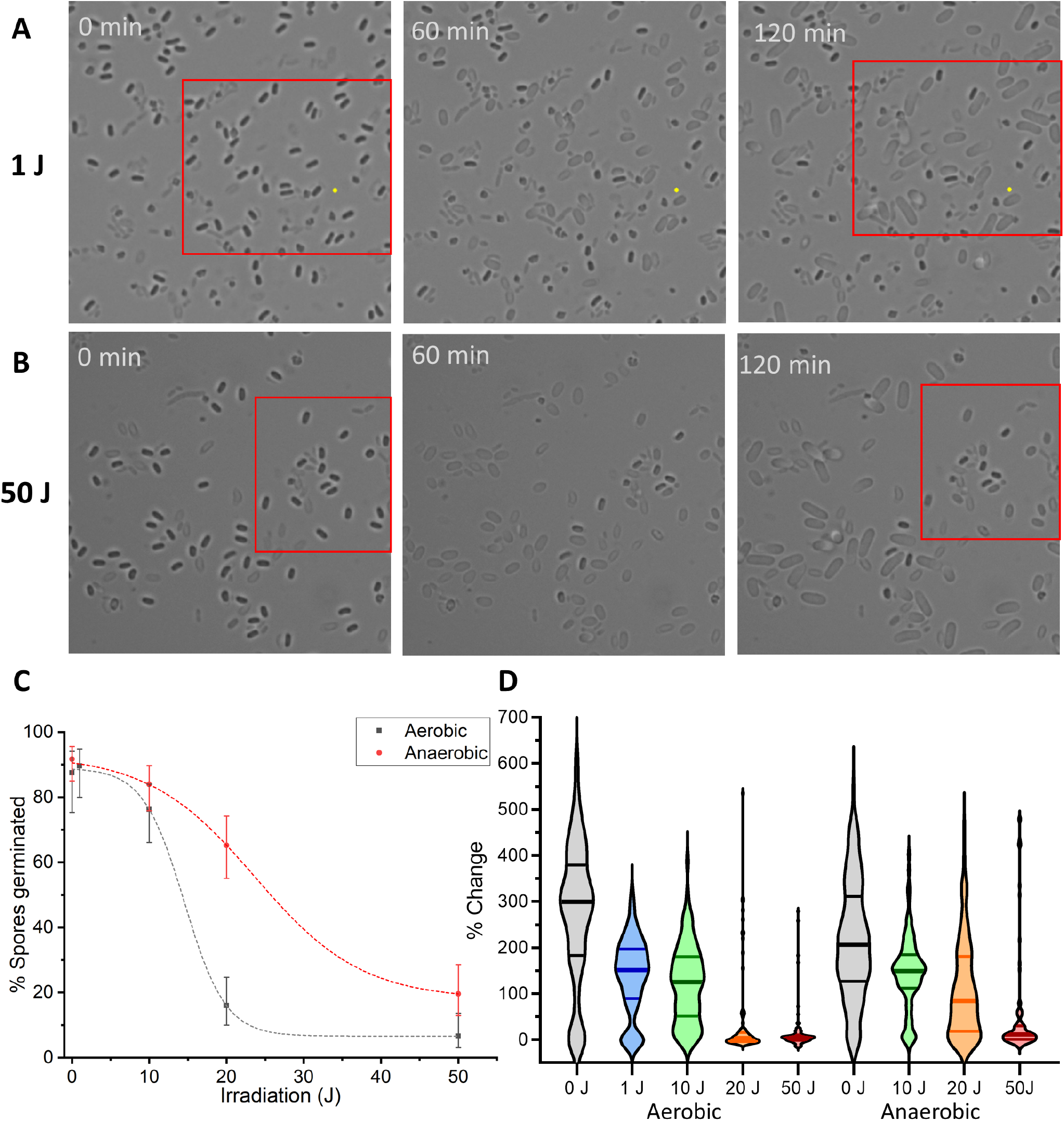
Representative field of view brightfield images of a time series of spores incubated in TSB over 120 minutes. The spores outlined in the red rectangle have been exposed to 1 J (A) and 50 J (B) with a 1064 nm laser. The irradiated spores remain in spore form after 120 minutes with 50 J irradiation but are unaffected by 1 J irradiation. Most non-irradiated spores germinate and grow into vegetative cells. (C) Differences in the rate of spore germination depends on spore irradiation and on the oxygen in the environment. Spore germination rate follows a sigmoid dose-response relationship vs. power used. Anaerobic incubation reduces phototoxic effects. (D) Violin plots show spores’ outgrowth (% change in size) into vegetative cells depending on irradition dose and incubation conditions.

We observe a Boltzmann sigmoid relation between the laser irradiation dose and percentage of spores germinated, see Figure 1 C. This is similar to classic dose-response curves described in literature ^38^. The sigmoid relationship indicates that spores can resist up to a few J of irradiation before losing the ability to germinate, but once the threshold is passed, the percentage of spores capable of surviving decreases rapidly. We believe the reason for this threshold may be spores’ defense against oxidative stress. Spores have several mechanisms to protect against ROS, including superoxide transmutases, small acid-soluble proteins and DNA repair mechanisms ^39,40^. However, the spore is metabolically inactive, and these mechanisms can be depleted, and once they are, generated ROS can damage the spores, as observed in our results.

The Boltzmann sigmoid relation of spore germination also relates well to our quantified assessment of spore size change. From brightfield images, we measured the size of cells after 2 h for different doses in aerobic media, see Figure 1 D. The change of spore size after exposure to doses of 20 J and 50 J is significantly different than the controls.

To visually assess if laser light exposure damaged the bacterial spore bodies we used SEM imaging. A SEM image of 50 J laser-irradiated spores, in which the TSB broth was washed off after treatment, is shown in Figure 2 A. As can be seen, there is a large variation in spore appearance, with some spores (orange arrows) appearing intact while others (blue arrows) are collapsed. Both the intact (with CaDPA) and collapsed (without CaDPA) spores did not germinate. This is consistent with a previous study using laser tweezers Raman spectroscopy which detected that CaDPA leaked out of spores after a high dose of laser irradiation ^10^. The results in this study thus confirm previous results and show that germination-inhibited spores do not necessarily need to appear collapsed. Compared to the non-germinating spores, non-irradiated spores (Figure 2 B) turn more elongated, with varying lengths as they are in different germination stages. This variation is expected since there is a high heterogeneity of spore germination rates for bacillus ^6^.

**FIGURE 2.**
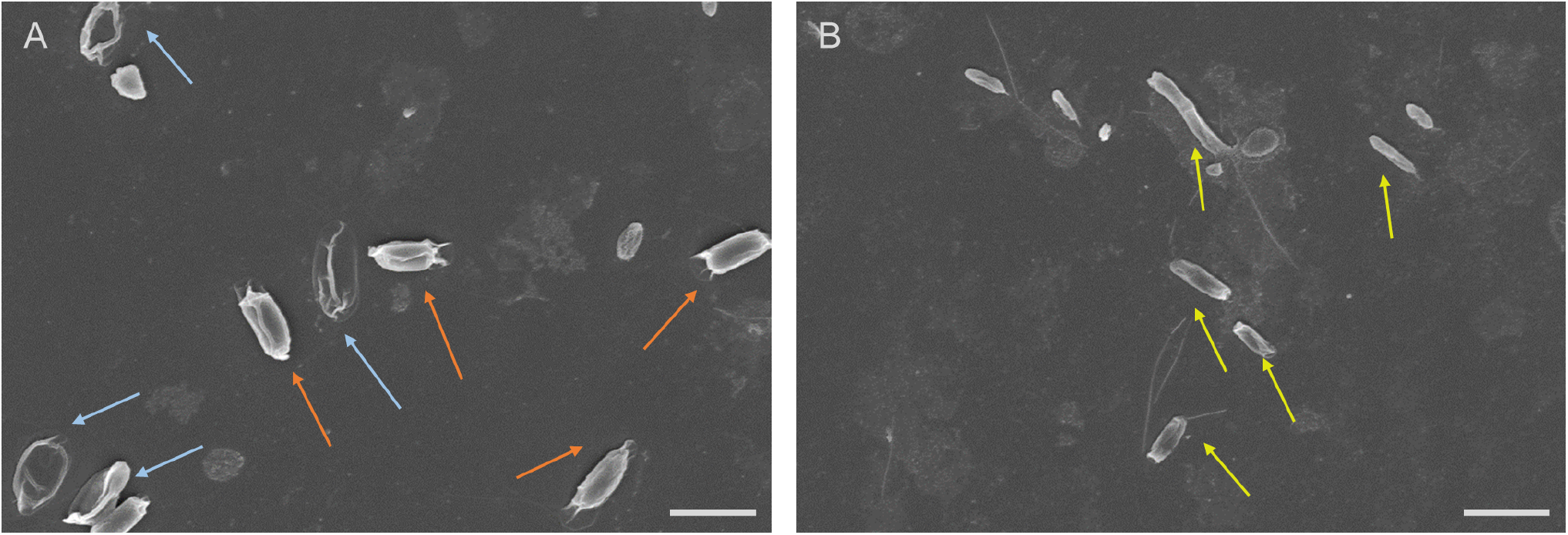
SEM images of spores incubated in TSB for 60 minutes. Both images are taken from the same sample with different fields of view. Spores that were irradiated with the 50 J (A) appear either intact (orange arrow) or collapsed (blue arrow), while spores unexposed to the laser (B) have started to germinate into vegetative cells (yellow arrows). Scale bars are 2 *µ*m.

We then tested whether ROS generation from molecular oxygen is the mechanism inhibiting germination as predicted. As discussed previously, the primary ROS generation mechanism reported in the literature for NIR lasers is singlet oxygen generation ^26^. While several possible strategies exist to limit the effect of ROS, such as using different ROS scavengers ^41^, these have disadvantages. The first disadvantage is that these ROS scavengers can have other chemical effects, with, for example, sodium azide being bactericidal and glucose oxidase decreasing the glucose content in growth media. The second is that these scavengers will not necessarily permeate inside the spore, where ROS may more effectively damage the spore.

A more direct method that can be used with facultative anaerobes such as *Bacillus thuringiensis* is to simply remove the oxygen from the growth media by degassing it in an anaerobic environment and then growing the cells anaerobically. The hypothesis is that anaerobic incubation would reduce the amount of oxygen within the spores, and thus reduce the effect of the laser beam in the optical tweezers on the spores. We found this hypothesis to be valid and the effect of the laser exposure on spores was significantly smaller on spores in an anaerobic environment (Figure 1 C, Figure S4 - S6). We saw that 92 % (n=109) of the non-irradiated spores germinated in anaerobic conditions, similar percentage as in aerobic conditions. Of the irradiated spores, 84 % (n=93) of the spores germinated with 10 J, 65 % (n=92) with 20 J, and 19 % (n=97) with 50 J. The germination of spores exposed to 10 J was not significantly different from control (p = 0.99), while germination of spores exposed to 20 J and 50 J was significantly below control (p = 0.005 and p < 0.0001 respectively) but significantly above the germination rate of spores incubated in an aerobic environment (p < 0.0001 and p = 0.01). The results thereby indicate that molecular oxygen is a major contributor to the germination-inhibiting effect we observe. However, other mechanisms still exist since there is still a measurable decrease in spore germination with 20 J, and at 50 J irradiation.

We have considered photothermal damage as a potential source of phototoxicity seen in anaerobic environments. However, based on published studies, temperature increases from laser power are expected to be small. We previously created a multi-physics simulation model for spore irradiation by a laser beam ^24^, and after adapting it to the new laser power and wavelength, the temperature increase of the spore content from a 1 W 1064 nm laser was calculated to be only 2.4 °C. Such a temperature increase would not affect the germination ability of spores, which can tolerate temperatures *>* 100 °C, nor would it be expected to cause mechanical damage from thermal expansion.

At high laser intensities, multi-photon absorption can also be a non-linear ROS generation source. Multi-photon absorption can create effects equivalent to visible and UV intense irradiation and lead to cell death ^42^ At higher laser powers the intensity in optical tweezers can amount to several MW/mm^2^, at this intensity multi-photon absorption is possible. Multi-photon absorption can generate ROS and cause photodamage independently from molecular oxygen, so that it can account for the smaller but still statistically significant reduction in spore germination for 20 J anaerobic experiments. Testing whether this is the case would require a long-time low-power exposure, however this type of assay would be complicated to perform since spores may begin to germinate before the irradiation process is completed.

## 4 CONCLUSION

Optical tweezers are a very useful tool for characerizing biophysical and physicochemical properties of small biological objects such as cells, bacteria, and spores. However, phototoxicity due to ROS production from optical tweezers must be accounted for during experiments. Typically spores are assumed to be highly resilient to environmental damage. However, we show that phototoxicity from 1064 nm optical tweezers can decrease the viability of spores, with irradiation above 10 J significantly suppressing their ability to germinate. This germination suppression follows a dose-response sigmoid relationship, indicating depletion of the spore’s mechanisms to counter oxidative stress. We further show that this effect is in large part driven by the molecular oxygen dissolved in water. The germination-inhibiting effect from a laser is reduced when spores are incubated in anaerobic broth, in line with theory prediction that ROS are generated from singlet oxygen generation. Overall, we hope this study highlights the risk for optical tweezers to unintentionally affect spores and ways to mitigate it.

## ^0^Abbreviations

ROS: reactive oxygen species

## 5 ACKNOWLEDGEMENTS

This work was supported by the Swedish Research Council (2019-04016); the Umeå University Industrial Doctoral School (IDS); Kempestiftelserna (JCK-1916.2). We thank Anna-Lena Johansson at FOI for providing the original stock of spores for this project.

The authors acknowledge the facilities and technical assistance of the Umeå Core Facility for Electron Microscopy (UCEM) at the Chemical Biological Centre (KBC), Umeå University, a part of the National Microscopy Infrastructure NMI (VR-RFI 2016-00968)

## References

1. Setlow P. Spores of Bacillus subtilis: Their resistance to and killing by radiation, heat and chemicals. Journal of Applied Microbiology 2006; 101(3): 514–525. doi: 10.1111/j.1365-2672.2005.02736.x

2. Stewart GC. The Exosporium Layer of Bacterial Spores: a Connection to the Environment and the Infected Host. Microbiology and Molecular Biology Reviews 2015; 79(4): 437–457. doi: 10.1128/MMBR.00050-15

3. Manchee RJ, Broster MG, Anderson IS, Henstridge RM, Melling J. Decontamination of Bacillus anthracis on Gruinard Island?. Nature 1983; 303(5914): 239–240. doi: 10.1038/303239a0

4. Goel AK. Anthrax: A disease of biowarfare and public health importance. World Journal of Clinical Cases 2015; 3(1): 20. doi: 10.12998/wjcc.v3.i1.20

5. Zhang P, Kong L, Setlow P, Li Yq. Characterization of Wet-Heat Inactivation of Single Spores of Bacillus Species by Dual-Trap Raman Spectroscopy and Elastic Light Scattering. Applied and Environmental Microbiology 2010; 76(6): 1796–1805. doi: 10.1128/AEM.02851-09

6. Zhang Y, Mathys A. Superdormant Spores as a Hurdle for Gentle Germination-Inactivation Based Spore Control Strategies. Frontiers in Microbiology 2019; 9(JAN): 1–10. doi: 10.3389/fmicb.2018.03163

7. Frentz Z, Dworkin J. Bioluminescence dynamics in single germinating bacterial spores reveal metabolic heterogeneity. Journal of The Royal Society Interface 2020; 17(170): 20200350. doi: 10.1098/rsif.2020.0350

8. Fällman E, Schedin S, Jass J, Andersson M, Uhlin BE, Axner O. Optical tweezers based force measurement system for quantitating binding interactions: system design and application for the study of bacterial adhesion. Biosensors and Bioelectronics 2004; 19(11): 1429–1437. doi: dx.doi.org/10.1016/j.bios.2003.12.029

9. Pesce G, Rusciano G, Sasso A, Isticato R, Sirec T, Ricca E. Surface charge and hydrodynamic coefficient measurements of Bacillus subtilis spore by optical tweezers.. Colloids and surfaces. B, Biointerfaces 2014; 116: 568–75. doi: 10.1016/j.colsurfb.2014.01.039

10. Malyshev D, Dahlberg T, Wiklund K, Andersson PO, Henriksson S, Andersson M. Mode of Action of Disinfection Chemicals on the Bacterial Spore Structure and Their Raman Spectra. Analytical Chemistry 2021; 93(6): 3146–3153. doi: 10.1021/acs.analchem.0c04519

11. Svoboda K, Block SM. Optical trapping of metallic Rayleigh particles.. Optics letters 1994; 19(13): 930–932. doi: 10.1364/OL.19.000930

12. Ayano S, Wakamoto Y, Yamashita S, Yasuda K. Quantitative measurement of damage caused by 1064-nm wavelength optical trapping of Escherichia coli cells using on-chip single cell cultivation system. Biochemical and Biophysical Research Communications 2006; 350(3): 678–684. doi: 10.1016/j.bbrc.2006.09.115

13. Mirsaidov U, Timp W, Timp K, Mir M, Matsudaira P, Timp G. Optimal optical trap for bacterial viability. Physical Review E 2008; 78(2): 021910. doi: 10.1103/PhysRevE.78.021910

14. Zhang Y, Miao Z, Huang X, Wang X, Liu J, Wang G. Laser Tweezers Raman Spectroscopy (LTRS) to Detect Effects of Chlorine Dioxide on Individual Nosema bombycis Spores. Applied Spectroscopy 2019; 73(7): 774–780. doi: 10.1177/0003702818817522

15. Mohanty SK, Rapp A, Monajembashi S, Gupta PK, Greulich KO. Comet assay measurements of DNA damage in cells by laser microbeams and trapping beams with wavelengths spanning a range of 308 nm to 1064 nm. Radiation Research 2002; 157(4): 378–385. doi: 10.1667/0033-7587(2002)157[0378:CAMODD]2.0.CO;2

16. Jockusch S, Turro NJ, Thompson EK, Gouterman M, Callis JB, Khalil GE. Singlet molecular oxygen by direct excitation. Photochem. Photobiol. Sci. 2008; 7(2): 235–239. doi: 10.1039/B714286B

17. Bagrov IV, Belousova IM, Kiselev VM, Kislyakov IM, Sosnov EN. Observation of the luminescence of singlet oxygen at λ = 1270 nm under LED irradiation of CCl4. Optics and Spectroscopy 2012; 113(1): 57–62. doi: 10.1134/S0030400X1207003X

18. Mohanty SK, Sharma M, Gupta PK. Generation of ROS in cells on exposure to CW and pulsed near-infrared laser tweezers. Photochem. Photobiol. Sci. 2006; 5(1): 134–139. doi: 10.1039/B506061C

19. Chan JW, Esposito AP, Talley CE, Hollars CW, Lane SM, Huser T. Reagentless Identification of Single Bacterial Spores in Aqueous Solution by Confocal Laser Tweezers Raman Spectroscopy. Analytical Chemistry 2004; 76(3): 599–603. doi: 10.1021/ac0350155

20. Huang Ss, Chen D, Pelczar PL, Vepachedu VR, Setlow P, Li Yq. Levels of Ca 2+ -Dipicolinic Acid in Individual Bacillus Spores Determined Using Microfluidic Raman Tweezers. Journal of Bacteriology 2007; 189(13): 4681–4687. doi: 10.1128/JB.00282-07

21. Zhang P, Setlow P, Li Y. Characterization of single heat-activated Bacillus spores using laser tweezers Raman spectroscopy. Optics Express 2009; 17(19): 16480. doi: 10.1364/OE.17.016480

22. Kong L, Setlow P, Li Yq. Observation of the dynamic germination of single bacterial spores using rapid Raman imaging. Journal of Biomedical Optics 2013; 19(1): 011003. doi: 10.1117/1.jbo.19.1.011003

23. Wang S, Yu J, Suvira M, Setlow P, Li Yq. Uptake of and Resistance to the Antibiotic Berberine by Individual Dormant, Germinating and Outgrowing Bacillus Spores as Monitored by Laser Tweezers Raman Spectroscopy. PLOS ONE 2015; 10(12): e0144183. doi: 10.1371/journal.pone.0144183

24. Malyshev D, Öberg R, Dahlberg T, et al. Laser induced degradation of bacterial spores during micro-Raman spectroscopy. Spectrochimica Acta Part A: Molecular and Biomolecular Spectroscopy 2022; 265: 120381. doi: 10.1016/j.saa.2021.120381

25. Malyshev D, Öberg R, Landström L, Andersson PO, Dahlberg T, Andersson M. pH-induced changes in Raman, UV–vis absorbance, and fluorescence spectra of dipicolinic acid (DPA). Spectrochimica Acta Part A: Molecular and Biomolecular Spectroscopy 2022; 271: 120869. doi: 10.1016/j.saa.2022.120869

26. Blázquez-Castro A. Optical Tweezers: Phototoxicity and Thermal Stress in Cells and Biomolecules. Micromachines 2019; 10(8): 507. doi: 10.3390/mi10080507

27. Stangner T, Dahlberg T, Svenmarker P, et al. Cooke–Triplet tweezers: more compact, robust, and efficient optical tweezers. Optics Letters 2018; 43(9): 1990. doi: 10.1364/OL.43.001990

28. Dahlberg T, Malyshev D, Andersson PO, Andersson M. Biophysical fingerprinting of single bacterial spores using laser Raman optical tweezers. In: Guicheteau JA, Howle CR., eds. Proc. SPIE 11416, Chemical, Biological, Radiological, Nuclear, and Explosives (CBRNE) Sensing XXISPIE. ; 2020

29. Dahlberg T, Andersson M. Optical design for laser tweezers Raman spectroscopy setups for increased sensitivity and flexible spatial detection. Applied Optics 2021; 60(16): 4519. doi: 10.1364/AO.424595

30. Schindelin J, Arganda-Carreras I, Frise E, et al. Fiji: An open-source platform for biological-image analysis. Nature Methods 2012; 9(7): 676–682. doi: 10.1038/nmeth.2019

31. Escamez S, André D, Sztojka B, et al. Cell Death in Cells Overlying Lateral Root Primordia Facilitates Organ Growth in Arabidopsis. Current Biology 2020; 30(3): 455–464.e7. doi: 10.1016/j.cub.2019.11.078

32. Pryor WA, Houk KN, Foote CS, et al. Free radical biology and medicine: it’s a gas, man!. American Journal of Physiology-Regulatory, Integrative and Comparative Physiology 2006; 291(3): R491–R511. doi: 10.1152/ajpregu.00614.2005

33. Maynard S, Schurman SH, Harboe C, Souza-Pinto dNC, Bohr VA. Base excision repair of oxidative DNA damage and association with cancer and aging. Carcinogenesis 2008; 30(1): 2–10. doi: 10.1093/carcin/bgn250

34. Cadet J, Wagner JR. DNA Base Damage by Reactive Oxygen Species, Oxidizing Agents, and UV Radiation. Cold Spring Harbor Perspectives in Biology 2013; 5(2): a012559–a012559. doi: 10.1101/cshperspect.a012559

35. Gassman NR, Wilson SH. Micro-irradiation tools to visualize base excision repair and single-strand break repair. DNA Repair 2015; 31: 52–63. doi: 10.1016/j.dnarep.2015.05.001

36. Setlow P. Spore Resistance Properties. In: Washington, DC, USA: ASM Press. 2016 (pp. 201–215)

37. Sunde EP, Setlow P, Hederstedt L, Halle B. The physical state of water in bacterial spores. Proceedings of the National Academy of Sciences 2009; 106(46): 19334–19339. doi: 10.1073/pnas.0908712106

38. Brooun A, Liu S, Lewis K. A Dose-Response Study of Antibiotic Resistance in Pseudomonas aeruginosa Biofilms. Antimicrobial Agents and Chemotherapy 2000; 44(3): 640–646. doi: 10.1128/AAC.44.3.640-646.2000

39. Moeller R, Raguse M, Reitz G, et al. Resistance of Bacillus subtilis Spore DNA to Lethal Ionizing Radiation Damage Relies Primarily on Spore Core Components and DNA Repair, with Minor Effects of Oxygen Radical Detoxification. Applied and Environmental Microbiology 2014; 80(1): 104–109. doi: 10.1128/AEM.03136-13

40. Cybulski RJ, Sanz P, Alem F, Stibitz S, Bull RL, O’Brien AD. Four Superoxide Dismutases Contribute to Bacillus anthracis Virulence and Provide Spores with Redundant Protection from Oxidative Stress. Infection and Immunity 2009; 77(1): 274–285. doi: 10.1128/IAI.00515-08

41. Huang YH, Chung KL, Yang WN, Chiu SH. Efficient symmetry-based screening strategy to speed up randomized circle-detection. Pattern Recognition Letters 2012; 33(16): 2071–2076. doi: 10.1016/j.patrec.2012.06.016

42. König K, So PTC, Mantulin WW, Gratton E. Cellular response to near-infrared femtosecond laser pulses in two-photon microscopes. Optics Letters 1997; 22(2): 135. doi: 10.1364/OL.22.000135

