## Supporting information for "Reactive oxygen species generated by infrared laser light in optical tweezers inhibits the germination of bacterial spores"

### Optical tweezer-generated ROS suppression of bacterial spore germination

**Method for processing images in ImageJ**

- The image was duplicated ('Image' → 'Duplicate')
- The duplicated image was converted to greyscale ('Image' → 'Type' → '8-bit')
- The duplicated image was converted to binary ('Process' → 'Binary' → 'Make Binary') and inverted so that spores/cells are white and background dark. ('Edit' → 'Invert').
- Areas not to be measured were manually removed from the binary duplicated image (using an area selection tool and then 'Edit' → 'Clear')
- In some cases, the threshold needed to be adjusted to ensure that the software accurately pinpointed the areas to be measured ('Image' → 'Adjust' → 'Threshold')
- The measurements were set to include both area and intensity ('Analyze' → 'Set Measurements' and then ticking both the 'Area' and 'Mean Grey Value' boxes)
- The measurements were set to redirect from the duplicated image to the original ('Analyze' → 'Set Measurements' → 'Redirect to')
- The measurements were carried out ('Analyze' → 'Analyze particles'), selecting the option "Show outlines" to track the same spores at the 0 and the 120 min timepoints.

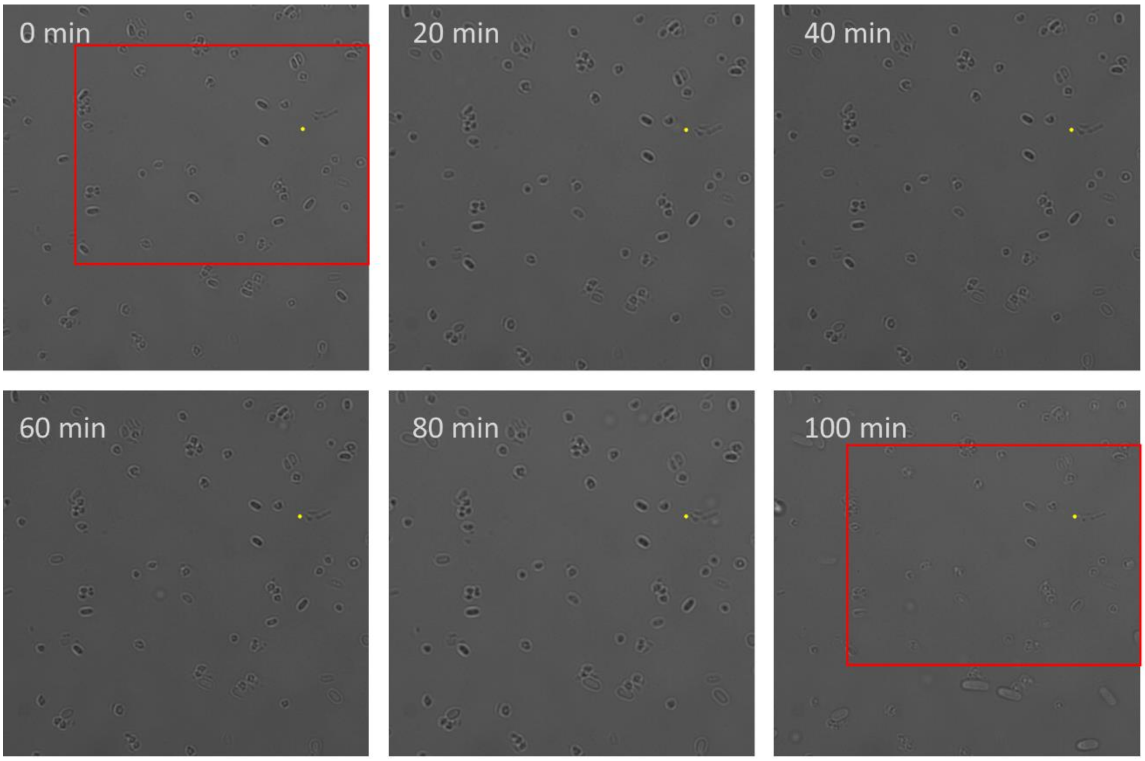


Figure S1. Representative example of a field of view of spores incubated in TSB over 120 minutes. The spores outlined in the red rectangle have been exposed to 20 J with a 1064 nm laser.

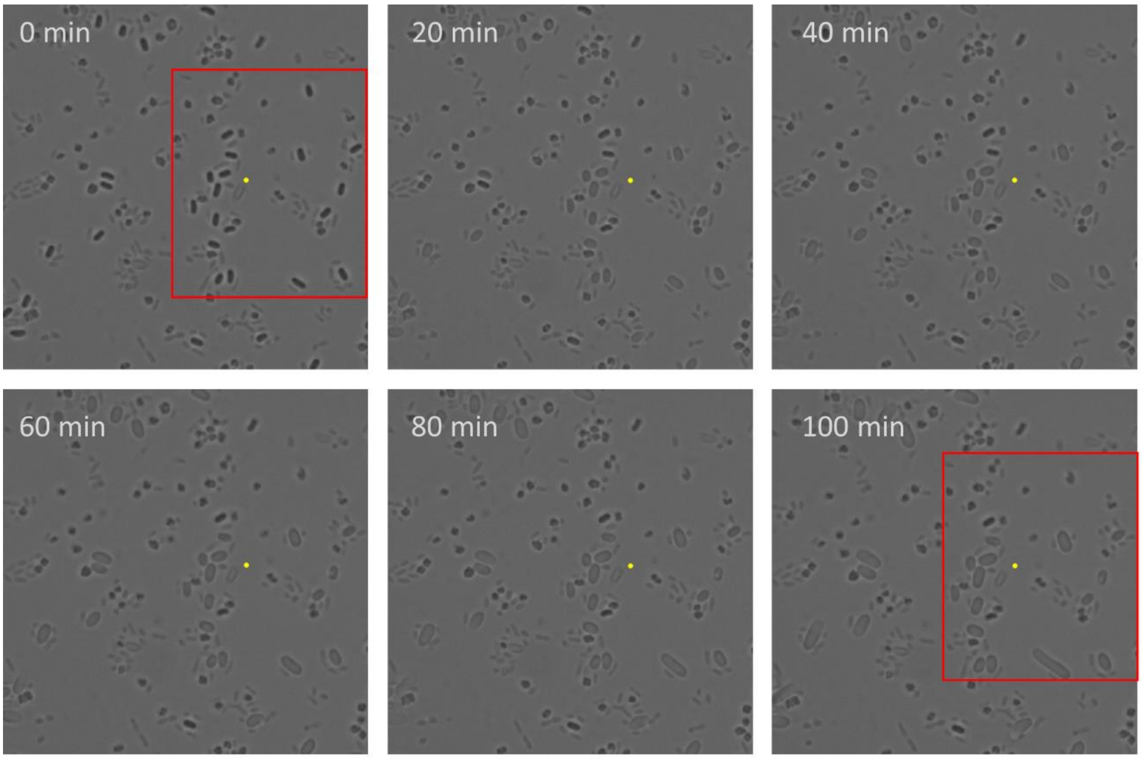


Figure S2. Representative example of a field of view of spores incubated in TSB over 120 minutes. The spores outlined in the red rectangle have been exposed to 10 J with a 1064 nm laser.

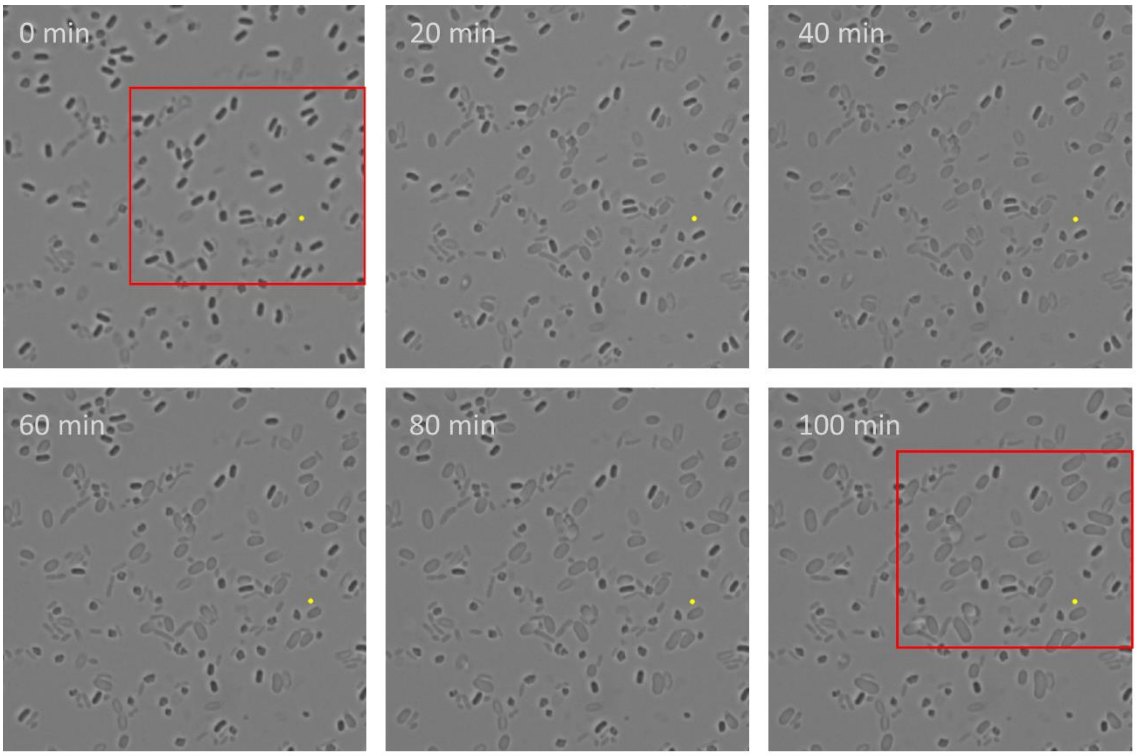


Figure S3. Representative example of a field of view of spores incubated in TSB over 120 minutes. The spores outlined in the red rectangle have been exposed to 1 J with a 1064 nm laser.

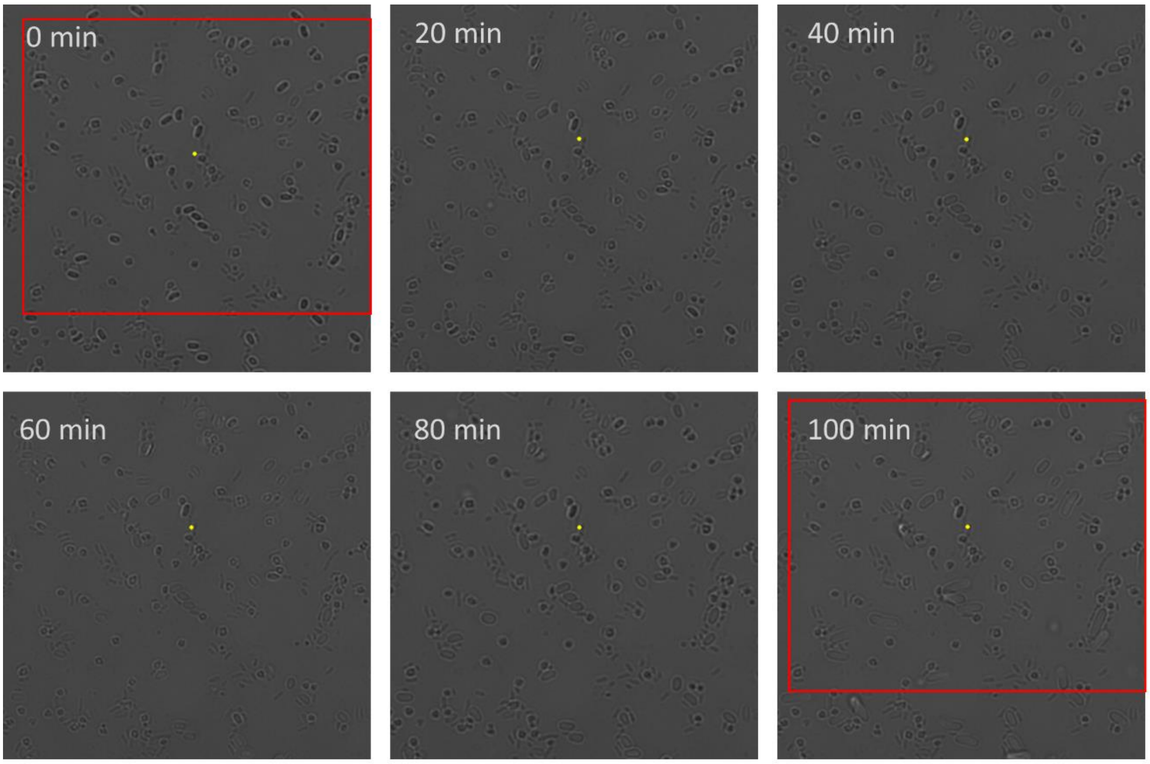


Figure S4. Representative example of a field of view of spores incubated in anaerobic TSB over 120 minutes. The spores outlined in the red rectangle have been exposed to 20 J with a 1064 nm laser.

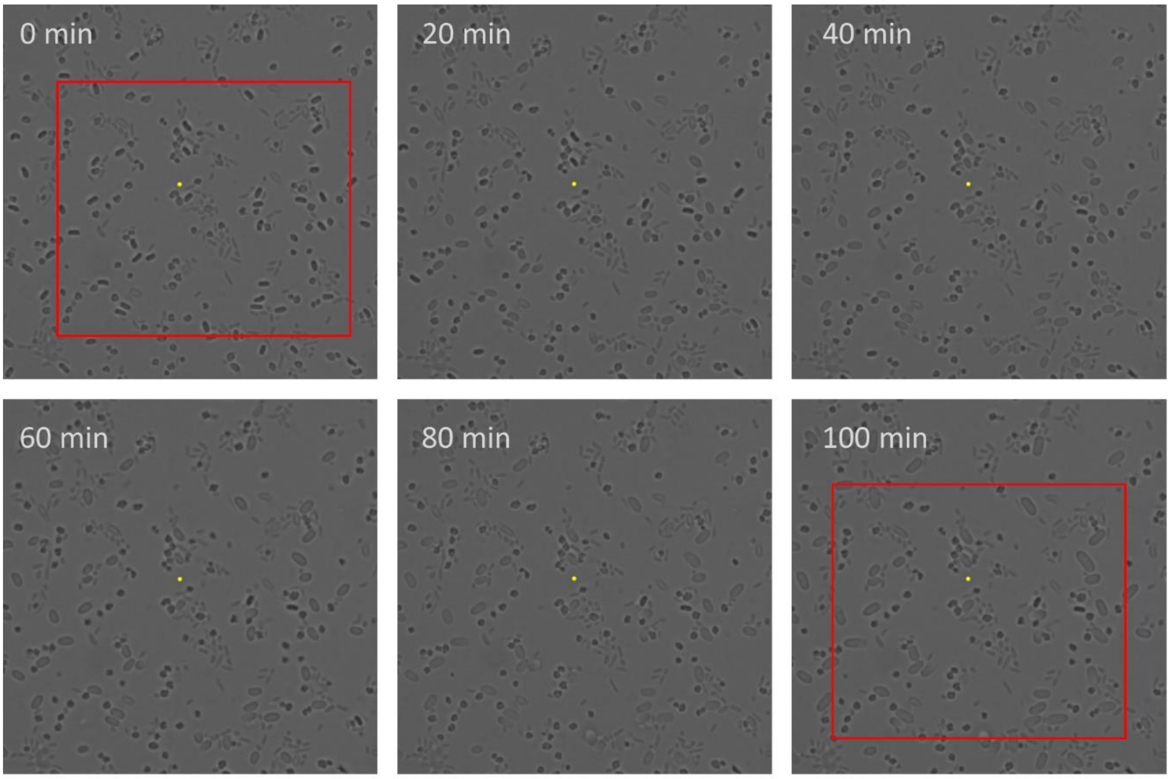


Figure S5. Representative example of a field of view of spores incubated in TSB over 120 minutes. The spores outlined in the red rectangle have been exposed to 10 J with a 1064 nm laser.

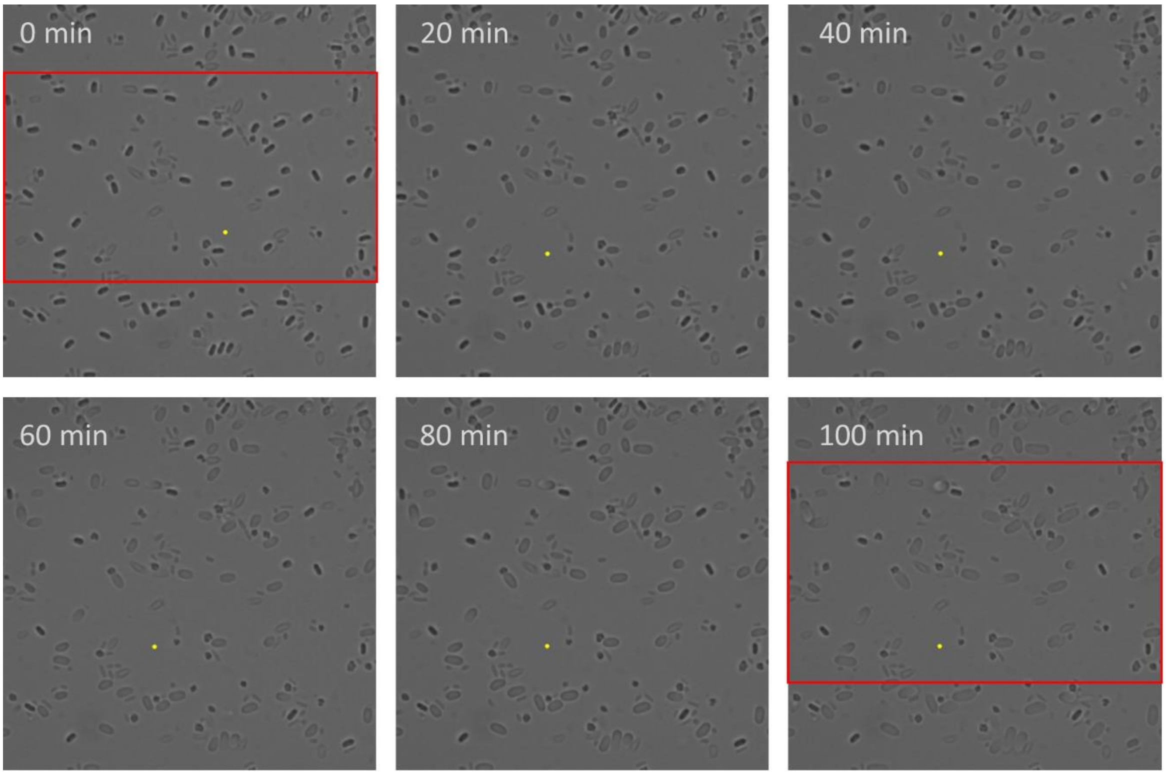


Figure S6. Representative example of a field of view of spores incubated in anaerobic TSB over 120 minutes. The spores outlined in the red rectangle have been exposed to 1 J with a 1064 nm laser.
